# Mitochondrial metabolism is rapidly re-activated in mature neutrophils to support stimulation-induced response

**DOI:** 10.1101/2025.02.03.636312

**Authors:** Jorgo Lika, James A. Votava, Rupsa Datta, Aleksandr M. Kralovec, Frances M. Smith, Anna Huttenlocher, Melissa C. Skala, Jing Fan

## Abstract

Neutrophils are highly abundant innate immune cells that are constantly produced from myeloid progenitors in the bone marrow. Differentiated neutrophils can perform an arsenal of effector functions critical for host defense. This study aims to quantitatively understand neutrophil mitochondrial metabolism throughout differentiation and activation, and to elucidate the impact of mitochondrial metabolism on neutrophil functions. To study metabolic remodeling throughout neutrophil differentiation, murine ER-Hoxb8 myeloid progenitor-derived neutrophils and human induced pluripotent stem cell-derived neutrophils were assessed as models. To study the metabolic remodeling upon neutrophil activation, differentiated ER-Hoxb8 neutrophils and primary human neutrophils were activated with various stimuli, including ionomycin, MSU crystals, and PMA. Characterization of cellular metabolism by isotopic tracing, extracellular flux analysis, metabolomics, and fluorescence-lifetime imaging microscopy revealed dynamic changes in mitochondrial metabolism. As neutrophils mature, mitochondrial metabolism decreases drastically, energy production is fully offloaded from oxidative phosphorylation, and glucose oxidation through TCA cycle is substantially reduced. Nonetheless, mature neutrophils retain the capacity for mitochondrial metabolism. Upon stimulation with certain stimuli, TCA cycle is rapidly activated. Mitochondrial pyruvate carrier inhibitors reduce this re-activation of the TCA cycle and inhibit the release of neutrophil extracellular traps. Mitochondrial metabolism also impacts neutrophil redox status, migration, and apoptosis without significantly changing overall bioenergetics. Together, these results demonstrate that mitochondrial metabolism is dynamically remodeled and plays a significant role in neutrophil function and fate. Furthermore, these findings point to the therapeutic potential of mitochondrial pyruvate carrier inhibitors in a range of conditions where dysregulated neutrophil response drives inflammation and contributes to pathology.

## 1. Introduction

Neutrophils are innate immune cells characterized by their active turnover and fast immune response. We seek to understand the dynamic metabolic remodeling associated with their maturation and support their immune response. As the most abundant leukocyte in circulation with a short lifespan, neutrophils need to be constantly produced from progenitors through multiple stages [1–4]. In the bone marrow, hematopoietic stem cells give rise to the common myeloid progenitor (CMP), which then gives rise to the granulocyte monocyte progenitor (GMP) [5]. GMPs are highly proliferative and can mature into neutrophil-specific progenitors. Then, these neutrophil-specific progenitors mature into circulating neutrophils and gain their capacity for performing effector functions. While the morphological and transcriptional changes during neutrophil differentiation are better characterized, the metabolic remodeling during this process is less understood. To model neutrophil maturation *in vitro*, a few systems have been developed. Murine progenitors can be conditionally immortalized in the GMP state by the transduction of bone marrow cells with estrogen receptor fusions of HOXB8. In the presence of estrogen, HOXB8 remains active and maintains the cells in their progenitor state, allowing for proliferation. Upon removal of estrogen, the cells differentiate and mature into neutrophils and gain phagocytic and inflammatory functions [6]. Similarly, human induced pluripotent stem cells can be differentiated into neutrophil-committed myeloblasts (CD34+ and CD33+) and then their maturation into functional mature neutrophils can be studied *in vitro* [7]. These models not only provide simple and effective methods of studying neutrophil differentiation but also generate functional neutrophils for studying the metabolism-immune function connection.

Once differentiated, neutrophils perform crucial functions in innate immunity. In circulation and tissues, neutrophils rapidly respond to damage- or pathogen-associated molecular patterns with an arsenal of effector functions which promote pathogen clearance and initiate inflammatory responses. First, neutrophils must migrate towards sites of infection guided by chemokine gradients in a process called chemotaxis [8]. Then, when they reach the affected site, they can undergo a specialized form of cell death called NETosis. In this process, neutrophils release their nuclear contents and granules (which contain antimicrobial peptides, proteases, and redox enzymes like myeloid peroxidase [MPO]) to create a harsh microenvironment that kills and prevents the spread of pathogens [8–10]. NETs are highly inflammatory and while they play a critical role in protecting the body from infection, their dysregulation contributes to a host of diseases, including autoimmune diseases, allergy, infection, and cancer [11,12]. Therefore, understanding the regulation of neutrophil fate and functions can identify novel therapeutic targets for addressing many pathologies.

Metabolic remodeling has been shown to be critical in cell differentiation, functional transitions, and cell death processes in many systems [13–16]. Although neutrophils are conventionally thought to be mainly glycolytic (and thus the role of mitochondrial metabolism had been underestimated), recent studies have found that neutrophils have great metabolic flexibility in a variety of pathways including the pentose phosphate pathway, fatty acid metabolism, and mitochondrial respiration [12,17–20]. Thus, mitochondrial metabolism can display specific activities to support a specific activated state. Research also suggests there are important functional changes in the role of mitochondria throughout the neutrophil life cycle [21]. Therefore, we seek to quantitatively characterize mitochondrial metabolism throughout neutrophil maturation using ER-Hoxb8 myeloid progenitor cells derived-neutrophils and human iPSC derived-neutrophils. We then investigated how mitochondrial metabolism is remodeled in neutrophils upon different stimulations. Based on these findings, we further elucidated the impact of mitochondrial metabolism remodeling on neutrophil cellular physiology and effector functions.

## 2. Results

### 2.1 Mitochondrial metabolism is decreased during neutrophil differentiation

To investigate the remodeling of mitochondrial metabolism throughout neutrophil differentiation, we first performed metabolomics over the time courses of two cell models, murine ER-Hoxb8 conditionally immortalized myeloid progenitor derived neutrophils (ER-Hoxb8 neutrophils), and human induced pluripotent stem cell derived neutrophils (iNeutrophils). In ER-Hoxb8 neutrophils, day 0 of differentiation (D0) corresponds with the GMP progenitor stage. Over the differentiation time courses, expression of mature neutrophil markers gradually increased, confirming the differentiation into neutrophils (Fig 1A) [6]. In iNeutrophils, day 0 corresponds to the myeloblast stage (CD34+ and CD33+). By day 7, they differentiate into neutrophil-like cells with increased expression of neutrophil markers including CD11b, CD15, CD16, CD66b, MPO, and lactoferrin, and gain effector functions as described in previous publications [22–24]. As ER-Hoxb8 neutrophils and iNeutrophils differentiate and mature, all reliably detected TCA cycle intermediates decreased substantially over time (Fig 1B-C). To further examine the changes in metabolic activity through the TCA cycle, we applied U^13^C-glucose tracing and measured the incorporation of glucose-derived carbon into the TCA cycle in undifferentiated (D0) and 5-day differentiated (D5) ER-Hoxb8 neutrophils. While labeled glucose contributed equally to glycolytic intermediates (i.e., 3-phosphoglycerate [3PG]) in undifferentiated and differentiated cells, label incorporation into TCA cycle intermediates was greatly reduced in differentiated cells (Fig 1D), suggesting a loss of mitochondrial glucose oxidation. Moreover, we measured the basal oxygen consumption rate (OCR) and found it reduced by 10-fold upon 5-day differentiation in ER-Hoxb8 neutrophils (Fig 1E). This further confirms an overall decrease in the mitochondrial metabolic rate, which may not be limited to reduced glucose oxidation. These results are consistent with previous reports that hematopoietic stem cells are less sensitive to glycolysis inhibition [28]. Moreover, as neutrophils mature, the mitochondria become more rod-like and decrease in biomass and number [27,28].

**Fig 1.**
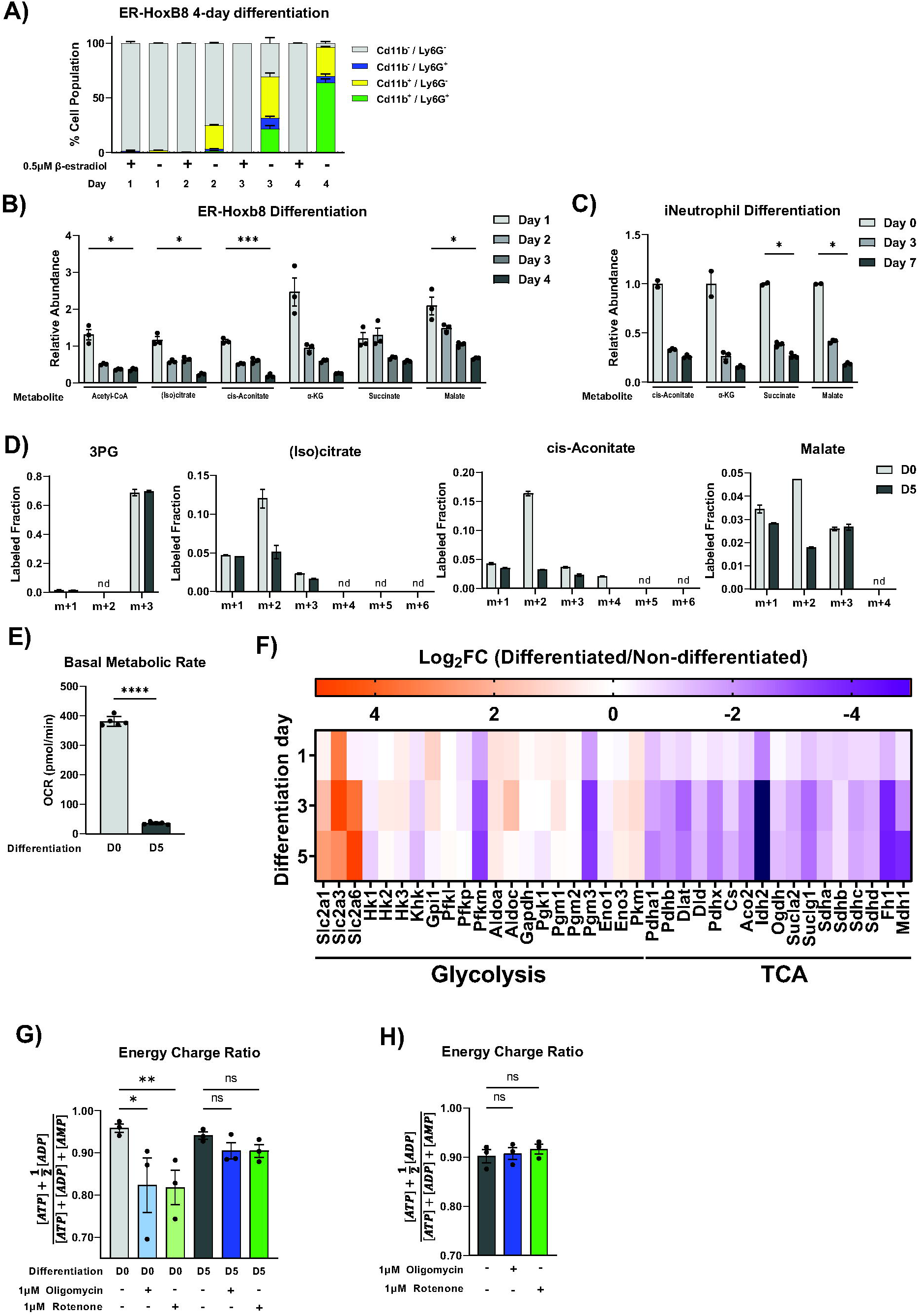
Mitochondrial metabolism is decreased during neutrophil differentiation. (A) Percentage of ER-Hoxb8 cells expressing neutrophil markers CD11b and Gr-1 throughout differentiation. Gating strategy is summarized in supplemental Figure S4. (B-C) Relative levels of TCA cycle metabolites across differentiation in ER-Hoxb8 neutrophils (B) and iNeutrophils (C). To determine significance, a mixed-effects model with Geisser-Greenhouse correction was run and p values were corrected for multiple comparisons. (D) Labeling of glycolysis and TCA intermediates from U^13^C-glucose in undifferentiated (D0) and 5-day differentiated (D5) ER-Hoxb8 neutrophils. Data representative of 3 independent experiments. (E) Oxygen consumption rate in undifferentiated (D0) vs differentiated (D5) ER-Hoxb8 neutrophils. Data are representative of 3 independent experiments. (F) Heatmap depicting expression of genes involved in glycolysis and the TCA cycle from a previously published RNA-seq dataset by Wang et al [22]. Day 5 differentiated ER-Hoxb8 neutrophils are compared to undifferentiated cells. (G) Energy charge ratio of undifferentiated (D0) vs differentiated (D5) ER-Hoxb8 neutrophils with or without 2h treatment of complex I (rotenone) or complex V (oligomycin) inhibitor. Data are combined from 3 independent experiments. Significance was determined with a Freidman test with Dunn’s multiple comparisons test. (H) Energy charge ratio measured in primary human neutrophils from 3 different donors with or without treatment of rotenone or oligomycin for 2 hours.

We further analyzed a previously published transcriptomic dataset of ER-Hoxb8 neutrophil differentiation [29] and found that many TCA cycle enzymes are transcriptionally downregulated, correlating with the metabolite level change (Fig 1F). To directly examine the contribution of mitochondrial metabolism to cellular bioenergetics, we treated undifferentiated and differentiated ER-Hoxb8 neutrophils with electron transport chain (ETC) inhibitors and measured cellular energy charge. ETC inhibitors significantly reduced cellular energy charge in undifferentiated ER-Hoxb8 neutrophils but had no significant impact in differentiated cells (Fig 1G). The independence of ETC is consistent with profoundly reduced TCA cycle activity upon neutrophil differentiation. However, despite such great reduction in mitochondrial metabolism, the base energy charge in differentiated cells is not significantly altered, indicating differentiated neutrophils can shift to the utilization of other pathways to fully maintain their bioenergetics. Finally, we examined the energy dependence of human peripheral blood neutrophils, which are mainly terminally differentiated mature neutrophils. Similar to differentiated ER-Hoxb8 neutrophils, ETC inhibitors have no impact on cellular energy levels of primary human peripheral blood neutrophils (Fig 1H).

### 2.2 Differentiated neutrophils reactivate TCA cycle metabolism upon certain stimulation

Although basal mitochondrial metabolism activity is greatly reduced in differentiated neutrophils, we found it can be rapidly re-activated upon stimulation. We performed metabolomics analysis in primary human neutrophils to examine the metabolomic changes upon activation with a variety of stimuli. Interestingly, pathway enrichment analysis of ionomycin stimulated neutrophils revealed the top significantly altered pathways are the malate-aspartate shuttle and the TCA cycle. Both pathways occur in the mitochondria (Fig 2A). Specifically, the top accumulated compounds upon ionomycin stimulation include most measured TCA cycle intermediates (increased by 10-fold or more), as well as some glycolytic intermediates (Fig 2A). U^13^C-glucose tracing in human peripheral blood neutrophils further showed that ionomycin stimulation significantly increased the rate of the incorporation of glucose into the TCA cycle, suggesting flux through the TCA cycle is substantially increased, which drives the accumulation of TCA intermediates (Fig 2B-C). At isotopic pseudo-steady state, the labeling of TCA cycle intermediates from glucose is minimal (<3%) in primary human neutrophils at baseline (Fig 2C). This is consistent with the oxygen consumption rate, which together suggests minimal base TCA cycle activity. Notably, this labeling fraction is rapidly increased by over 10-fold one hour after ionomycin stimulation. This degree of TCA cycle labeling increase greatly exceeds the increase of glucose’s contribution to glycolytic compounds (e.g. ∼2-fold increase in 3-labeled 3PG), demonstrating that neutrophil activation is coupled with a specific increase in mitochondrial oxidation beyond the general increase of glucose metabolism (which is reported in our previous paper [30]). For glucose to be oxidized by the TCA cycle, the glycolytic product, pyruvate, needs to be first transported into the mitochondria via the mitochondria pyruvate carrier (MPC). To further validate the specific increase in mitochondrial metabolism we treated primary human neutrophils with MPC inhibitors, UK-5099 and azemiglitazone. Treatment with either inhibitor completely inhibited the ionomycin-induced increase in label incorporation into the TCA cycle (Fig 2D-E). Notably, azemiglitazone treatment, but not UK-5099, also significantly increased the labeling in glycolysis intermediates (Fig 2E). This indicates that beyond its effect of inhibiting MPC, azemiglitazone has additional effects in increasing glycolysis, which has been suggested by previous literature [31]. Similar to human peripheral blood neutrophils, ionomycin stimulation of ER-Hoxb8 neutrophils also substantially increased label incorporation from glucose into the TCA cycle and this increase can be inhibited by UK-5099 (Fig 2F).

**Fig 2.**
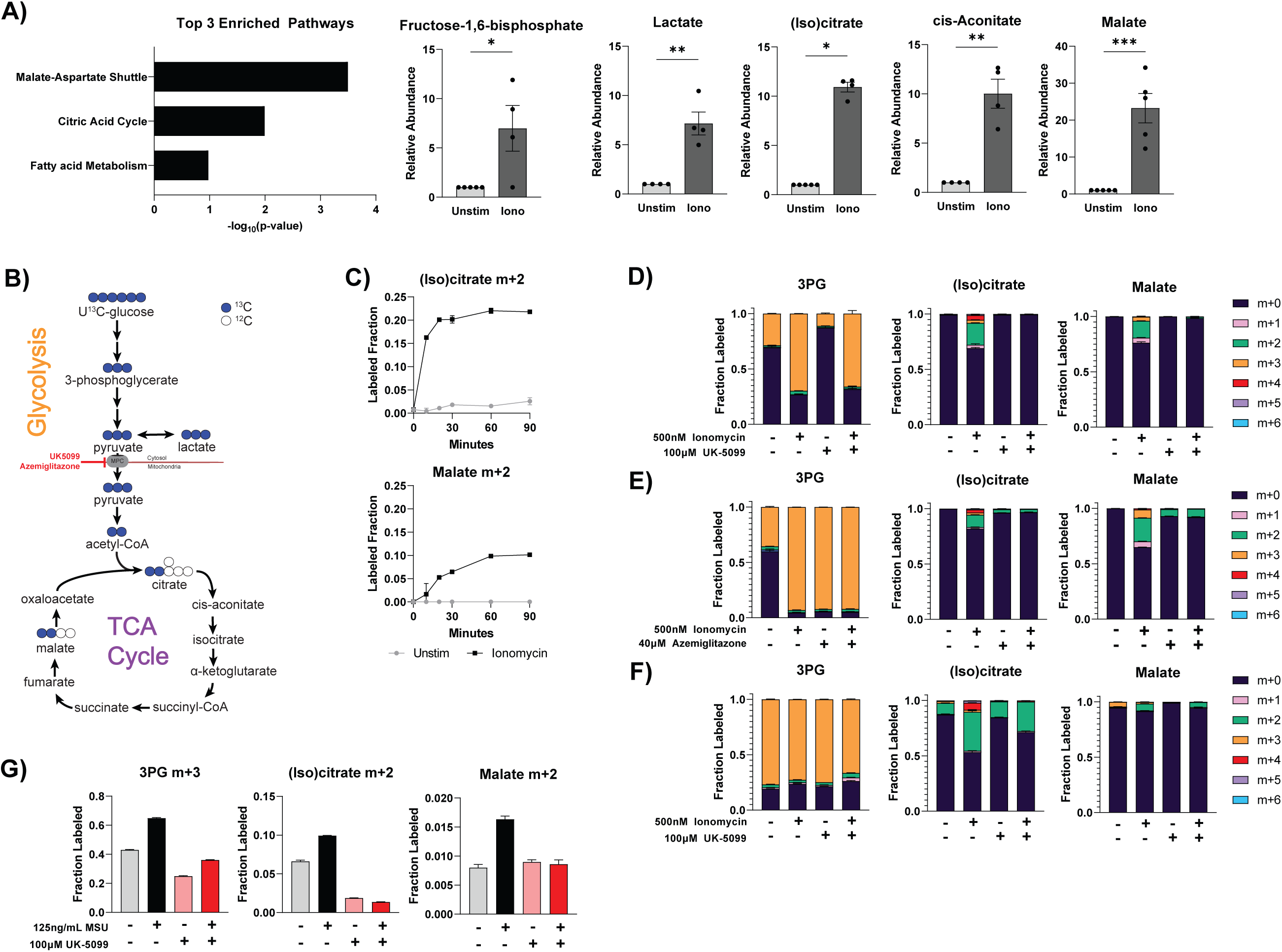
Differentiated neutrophils reactivate TCA cycle metabolism upon certain stimulation. (A) Pathway enrichment analysis on metabolomic changes induced by ionomycin stimulation and relative abundance of indicated metabolites in primary human neutrophils (n=5 different donors, individual dots present different biological replicates). Relative abundance of glycolysis and TCA metabolites was normalized by protein content and made relative to the unstimulated condition from the respective donor. Significance was determined by ratio paired t test with Holm-Šídák’s multiple comparisons test. (B) Schematic demonstrating labeling of glycolysis and TCA cycle from U^13^C-glucose. (C) Labeling fraction of indicated compound from U^13^C-glucose over time in primary human neutrophils. Data shown are representative of 2 different donors. (D-E) Labeling pattern of glycolysis and TCA intermediates after U^13^C-glucose labeling in primary human neutrophils treated under indicated conditions for 1 hour. Data shown are representative of experiments from at least 3 different human donors. (F) Labeling pattern of glycolysis and TCA intermediates after U^13^C-glucose labeling in differentiated ER-Hoxb8 neutrophils treated under specified conditions for 1 hour. Data are representative of at least two independent experiments. (G) Labeling of glycolysis and TCA intermediates from U^13^C-glucose in primary human neutrophils stimulated with or without monosodium urate (MSU) crystals under specified conditions for 1 hour. Data shown are representative of experiments from at least 3 different human donors.

Like ionomycin stimulation, activation with monosodium urate (MSU) crystals also led to significant increase in glucose-derived TCA cycle labeling, which can be blocked by the treatment of UK-5099 (Fig 2G). However, such a significant increase in TCA cycle labeling was not observed in phorbol myristate acetate (PMA) stimulation (Fig S1A). These results show that in mature neutrophils, although the utilization of the TCA cycle is minimal at baseline, the capacity of mitochondrial oxidation is retained, and TCA cycle is substantially and rapidly activated upon activation by specific stimuli.

### 2.3 Activation of mitochondrial glucose oxidation and effector functions are dependent on calcium availability

To investigate the molecular mechanisms that permit such rapidly increased TCA cycle flux, we examined the phosphorylation status of pyruvate dehydrogenase (PDH), which is the gate keeping enzyme of glucose oxidation in the mitochondria. In ER-Hoxb8 neutrophils, ionomycin stimulation led to rapid dephosphorylation of PDH, which is known to activate PDH (Fig 3A) [32]. Pyruvate dehydrogenase phosphatase_—_which dephosphorylates PDH—is activated by calcium [33]. Ionomycin is a calcium ionophore [34] and MSU crystal stimulation leads to calcium influx [35], whereas PMA has minimal changes in calcium influx compared to ionomycin [36]. Therefore, we hypothesized calcium signaling plays a vital role in the TCA cycle re-activation induced by these specific types of stimuli. To examine how calcium availability impacts neutrophil metabolism and functions, Ethylene glycol tetraacetic acid (EGTA) was used to reduce free extracellular calcium availability by chelating free calcium at a 1:1 ratio. EGTA treatment reduced glucose incorporation into TCA cycle intermediates, but not glycolysis intermediates, in a dose-dependent manner in ionomycin stimulated neutrophils (Fig 3B). Calcium concentration in RPMI is ∼400 µM, therefore EGTA at 500 µM completely chelates available calcium. At this level, there is minimal labeling incorporation into TCA cycle intermediates, like in unstimulated cells. This suggests activation of glucose mitochondrial oxidation is downstream of calcium signaling. EGTA also impaired ionomycin-induced NET release (Fig 3C-D). In imaging analysis, DNA release that exceeds the size of the cell (indicating true NETs versus lytic cell death), is lost with 500 µM EGTA (Fig 3C). Furthermore, ionomycin-induced MPO release was reduced by EGTA in a dose-dependent manner to baseline levels when EGTA is increased to 500 µM (Fig 3D). Lastly, complete chelation of calcium significantly reduced neutrophil migration (Fig 3E). Together, these data demonstrate that activation of mitochondrial glucose oxidation and neutrophil functions depend on calcium availability.

**Fig 3.**
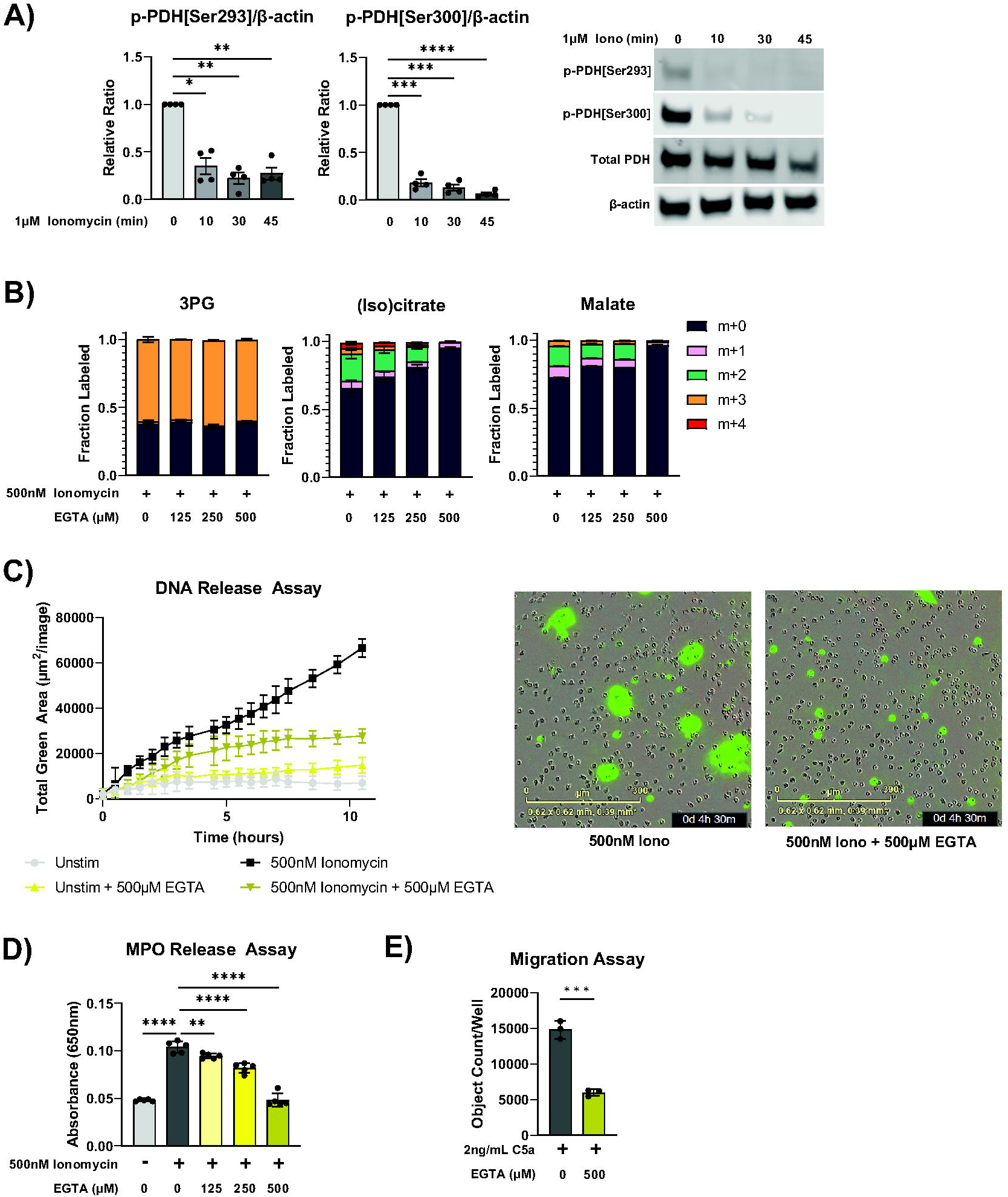
Mitochondrial metabolism and neutrophil response in varied calcium availability. (A) Immunoblot of phospho-PDH[ser293], phospho-PDH[ser300], total PDH E1 subunit, and β-actin in differentiated ER-Hoxb8 neutrophils stimulated with ionomycin for indicated duration. Relative signals (normalized by β-actin from the same blot) from 4 different independent experiments were made relative to the unstimulated condition and compiled. Representative images of immunoblots are shown to the right. Significance was determined with RM two-way ANOVA with Geisser-Greenhouse correction and Šídák’s multiple comparisons test. (B) Labeling pattern of glycolysis and TCA intermediates from U^13^C-glucose in primary human neutrophils treated under specified conditions for 1 hour. Data shown are representative of experiments from 3 different human donors. (C) Extracellular DNA release over time of primary human neutrophils cultured under specified conditions. Representative microscopy images at the 4.5-hour time point are shown to the right. Data shown are representative of experiments from 3 different human donors. (D) MPO activity assay from supernatant of primary human neutrophils treated for 4 hours under specified conditions. Data shown are representative of experiments from 3 different human donors. Significance was determined by ordinary one-way ANOVA and Šídák’s multiple comparisons test. (E) Number of neutrophils that migrated through a transwell to C5a after 1.5 hours under specified treatment conditions. Data shown are representative of experiments from 3 different human donors. Significance was determined by ordinary one-way ANOVA and Šídák’s multiple comparisons test.

### 2.4 Stimulated neutrophils adopt a different redox and energy state

Given the role of TCA cycle in redox and energy metabolism, we next examined how ionomycin stimulation and MPC inhibition altered energy charge and cellular redox state. Despite increased TCA cycle activity, ionomycin stimulation significantly decreases cellular energy charge (Fig 4A), suggesting a more overwhelming increase in energy consumption upon activation. UK-5099 and azemiglitazone do not significantly alter energy charge in the unstimulated or stimulated cells.

**Fig 4.**
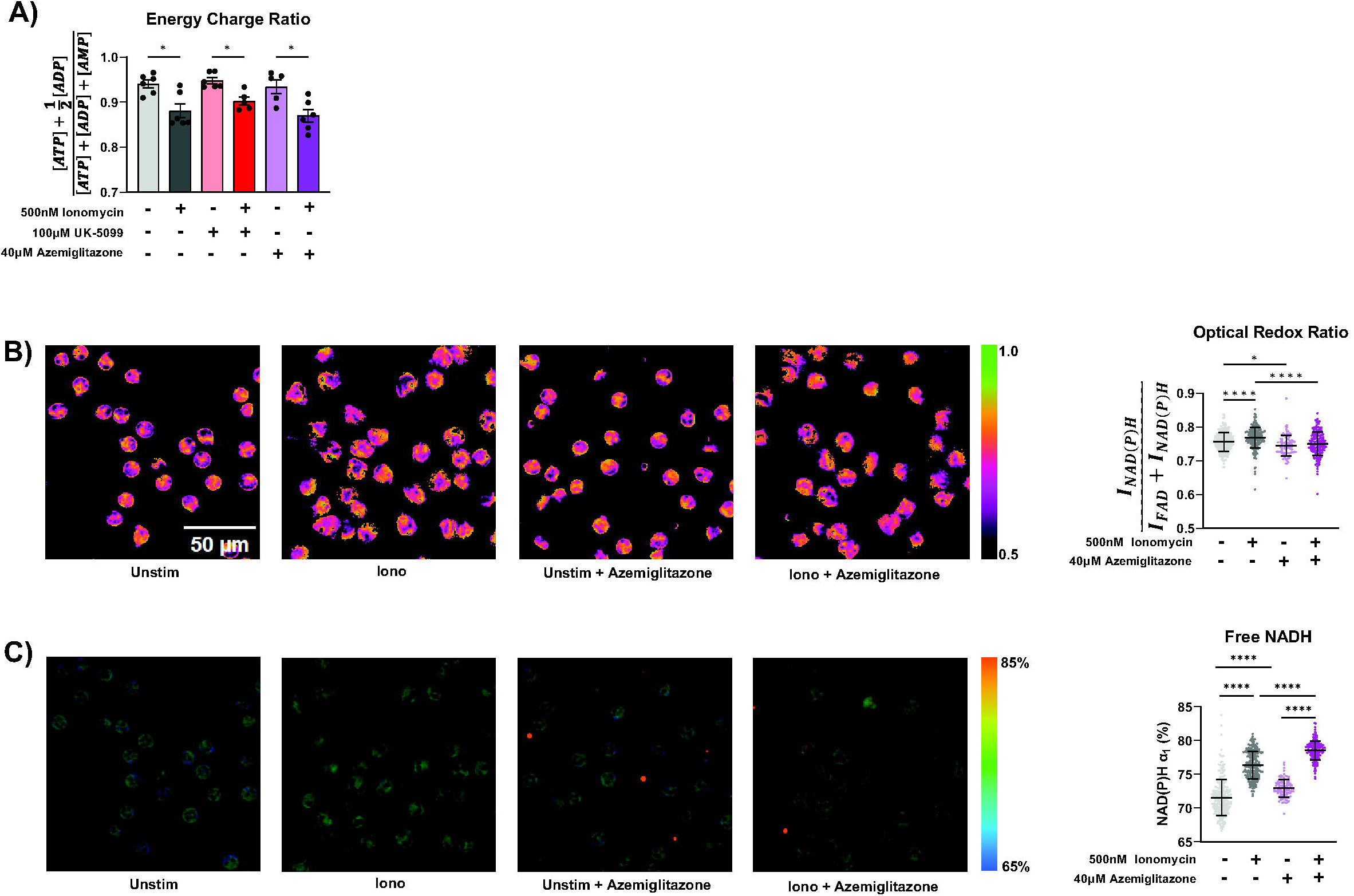
Redox and energy state in stimulated neutrophils. (A) Energy charge measured in primary human neutrophils from 6 different donors treated for 1 hour under specified conditions. Significance was determined with RM two-way ANOVA with Geisser-Greenhouse correction and Šídák’s multiple comparisons test. (B) Representative images of optical redox ratio [I_NAD(P)H_ / (_INAD(P)H_ + I_FAD_)] and corresponding single cell quantification (right) of neutrophils treated under specified conditions for 90 minutes. Significance was determined with ordinary one-way ANOVA and Tukey’s multiple comparisons test. (C) Representative images of free NAD(P)H percentage (α1) in neutrophils treated under specified conditions for 90 minutes and corresponding single cell quantification (right). Significance was determined with ordinary one-way ANOVA and Tukey’s multiple comparisons test. 5-6 images were acquired per condition, each data point is a single cell. There are at least 100 cells/condition.

To profile cellular redox status, we performed optical metabolic imaging using endogenous metabolic coenzymes NAD(P)H and FAD, which provides single cell optical redox ratio. Additionally, fluorescent lifetime imaging microscopy (FLIM) provides insights into protein binding of NAD(P)H. Optical metabolic imaging has recently been shown to be an effective method to characterize metabolic changes associated with activation in live neutrophils [37]. Upon ionomycin stimulation, there is a significant shift towards a more reduced redox state, as measured by the increase of the optical redox ratio (Fig 4B). This is likely due to increased TCA cycle activity producing NADH, which we consistently observed with LC-MS based metabolomics (Figure S2C), causing a reduction of FAD. The percentage of NAD(P)H in the free form (α1) is also significantly increased (Fig 4C). Azemiglitazone further increased the percentage of free NADH, in both the stimulated and unstimulated condition (Fig 4C). This is likely due to increased free NADH production from increases in glycolysis induced by azemiglitazone. However, azemiglitazone reduced the optical redox ratio, consistent with MPC inhibition. MPC inhibition would limit the utilization of multiple nutrients_—_such as glucose, glycogen, extracellular pyruvate—in the mitochondria, limiting TCA cycle activity and ultimately the reduction of FAD and NAD. This would prevent the stimulation-induced shift towards a more reduced state. Together, these results suggest the activation of the TCA cycle is uncoupled from overall bioenergetic changes but important for the shifts in cellular redox state upon ionomycin stimulation. It also further demonstrates the potential to use FLIM to characterize neutrophils in different activation states.

### 2.5 MPC inhibitors alter neutrophil fate and function

Next, we examined the impact of perturbing mitochondrial metabolism on stimulation induced neutrophil functions. Ionomycin is known to stimulate NADPH Oxidase (NOX)-independent neutrophil NET release [34]. Consistently, we observed an increase of extracellular DNA release over time after ionomycin stimulation. Strikingly, such ionomycin induced DNA-release is profoundly suppressed by both UK-5099 and azemiglitazone (Fig 5A-B). Similarly, DNA release induced by MSU crystals was also strongly inhibited by UK-5099 (Fig 5C). In contrast, UK-5099 had little effect on PMA-induced DNA release (Fig 5D) [38]. As release of NOX-independent NETs often requires histone citrullination to decondense chromatin, we further examined citrullinated histone levels and found UK-5099 treatment significantly reduced ionomycin-induced histone citrullination (Fig 5E). Ionomycin-induced MPO release was also profoundly reduced by MPC inhibitors (Fig 5F). These results demonstrate the important role of mitochondrial metabolism in NETosis induced by specific stimuli.

**Fig 5.**
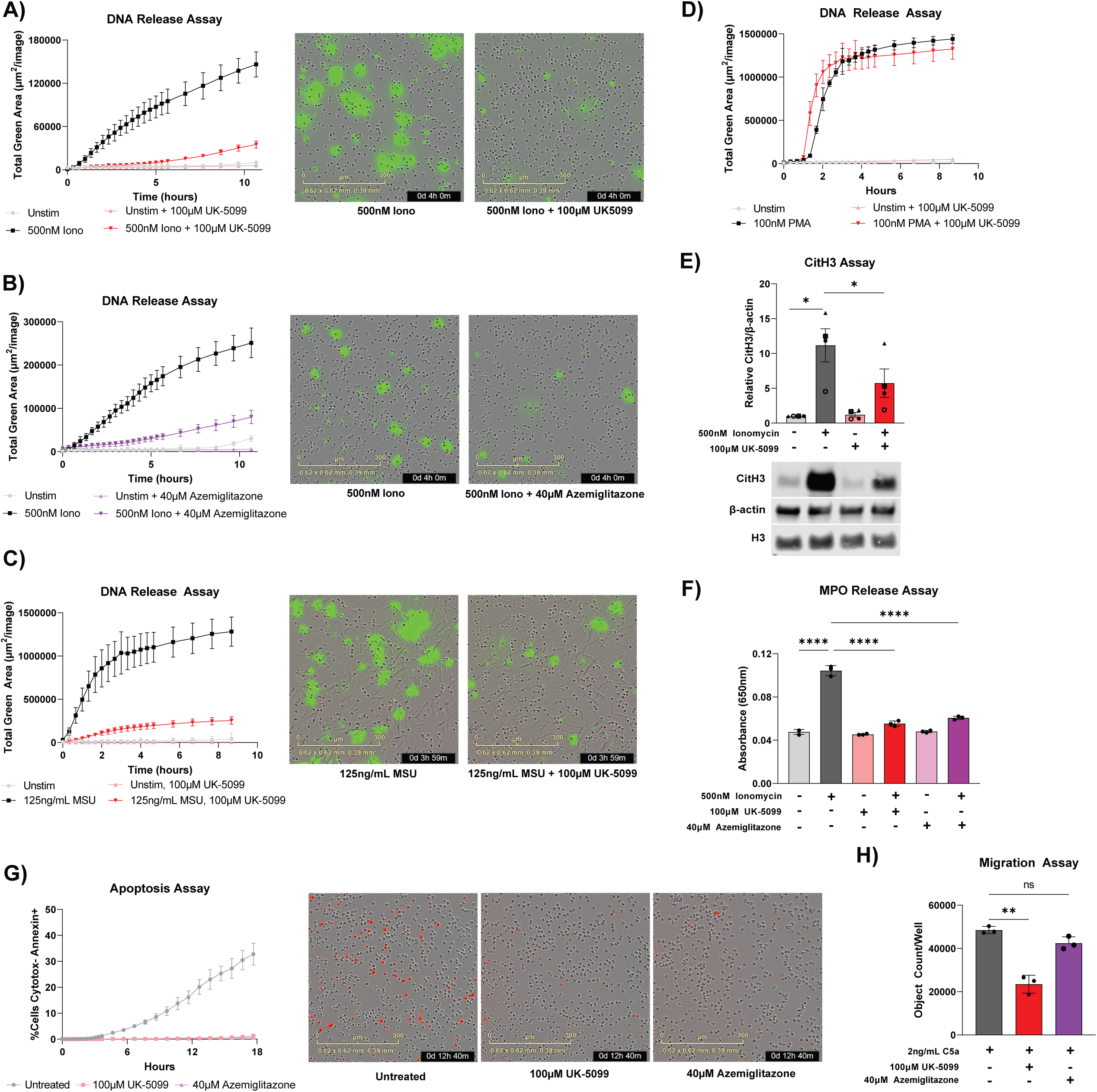
Mitochondrial pyruvate carrier (MPC) inhibitors alter neutrophil response and survival. (A-C) Extracellular DNA release over time of primary human neutrophils stimulated with ionomycin or MSU crystals, with or without treatment of MPC inhibitors and specified in legends. Data shown are representative from 3 different human donors and representative microscopy images at the 4-hour time point are shown to the right. (D) Extracellular DNA release over time of primary human neutrophils stimulated with PMA, with or without treatment of UK-5099. (E) Immunoblot of citrullinated histone 3 (CitH3), histone 3 (H3), and β-actin in cell lysate from primary human neutrophils treated for 4 hours under specified treatment conditions. Relative CitH3 signal (normalized by β-actin signal from the same blot) from 4 different donors were made relative to the unstimulated condition and compiled. Representative images of immunoblots are shown below. Significance was determined with RM one-way ANOVA with Geisser-Greenhouse correction and Šídák’s multiple comparisons test. (F) MPO activity assay from supernatant of primary human neutrophils treated for 4 hours under specified treatment conditions. Data shown are representative of experiments from 3 different human donors. Significance was determined by ordinary one-way ANOVA and Šídák’s multiple comparisons test. (G) Apoptosis over time of primary human neutrophils cultured under specified conditions. Data shown are representative from 3 different human donors with representative microscopy images at the 12-hour time point shown to the right. Red signal is from annexin V staining. (H) Number of neutrophils that migrated through a transwell to C5a after 1.5 hours under specified treatment conditions. Data shown are representative of experiments from 3 different human donors. Significance was determined by ordinary one-way ANOVA and Šídák’s multiple comparisons test.

As short lived innate immune cells, neutrophils often undergo programmed death. In addition to NETosis, which is a form of stimulation-induced, highly inflammatory neutrophil death associated with the release of NETs, neutrophils can also undergo apoptosis. In an unstimulated state, UK-5099 and azemiglitazone inhibited neutrophil apoptosis and extended neutrophil lifetime in *ex vivo* culture (Fig 5G).

Since calcium signaling and mitochondrial ATP production have also been shown to play a critical role in orienting the neutrophil during chemotaxis, we hypothesized that perturbing mitochondrial metabolism could also disrupt chemotaxis [39,40]. To examine the impact of MPC inhibitors on neutrophil migration capacity, we measured neutrophil migration towards chemoattractant in a transwell system. Migration is significantly reduced by UK-5099 treatment (Fig 5H). Together, these results demonstrate that MPC inhibitors can alter neutrophil functions.

## 3. Discussion

In this work, we quantitatively examined TCA cycle metabolism throughout neutrophil differentiation and upon neutrophil activation. The results reveal that mitochondrial metabolism is highly dynamic and plays a significant, context-specific role in neutrophils.

Mitochondrial metabolism is canonically considered to be an energy producing pathway. Therefore, its role in neutrophils, a highly glycolytic cell type with a low mitochondrial content, has been traditionally underappreciated. Indeed, our results agree with the previous reports that TCA cycle metabolism is strongly down regulated throughout neutrophil differentiation. Therefore, from a bioenergetic perspective, it has little contribution in mature neutrophils [21,41–43]. The highly consistent results on TCA cycle and ETC downregulation in ER-Hoxb8 neutrophils and iNeutrophils, as well as the lack of glucose-derived carbon in the TCA cycle in mature human peripheral blood neutrophils, supports that ER-Hoxb8 neutrophils and iNeutrophils are effective models to investigate metabolic rewiring across differentiation. This also highlights that the utilization of mitochondrial metabolism in neutrophils varies based on maturation state. Such maturation dependent difference has clinical relevance. Neutrophils in circulation can be a heterogeneous population. While the percentage of immature neutrophils is low in healthy blood, it is elevated in patients with cancer and other autoimmune diseases [44,45]. It has been shown in a cancer model that immature neutrophils use mitochondrial metabolism to support the production of ROS, which suppresses anti-tumor T-cell responses [46]. In these disease contexts, the metabolic heterogeneity between mature and immature neutrophils can contribute to the disease progression and is of interest for therapeutic targeting.

Even though mature neutrophils switch to ETC-independent energy production, we found mitochondrial capacity is maintained and can be greatly activated within minutes upon specific stimulation. Moreover, we demonstrated that treating neutrophils with inhibitors of MPC, a gatekeeper for carbon influx into TCA cycle, inhibited the stimulation-induced activation of the TCA cycle and substantially suppressed NET release and MPO release in activated neutrophils. Such strong functional effects of MPC inhibitors are not associated with any significant perturbation in energy charge, indicating the mechanism is uncoupled from overall cell bioenergetics. This supports the idea that in mature neutrophils mitochondria transition to be an organelle primarily important for signaling and regulatory functions. TCA cycle activation resulted in substantial changes in a series of important metabolites, including TCA cycle intermediates (such as citrate, cis-aconitate, and malate) and redox cofactors (such as NADH and FAD) (Fig 4 and S2). Current literature has demonstrated that many metabolites involved in the TCA cycle have immunoregulatory roles. For example, acetyl-CoA metabolism has been shown to regulate functions across various immune cell types through protein acetylation [47]. Succinate, α-ketoglutarate, and mitochondrial ROS were found to impact several signaling pathways, such as hypoxia inducible factor and nuclear factor-κB [48,49]. Understanding the specific mechanisms by which these MPC inhibitors regulate neutrophil functions is an important future direction.

The results here showed that neutrophils can rapidly and substantially remodel their mitochondrial metabolism upon activation. This further demonstrates the general idea—like our group and others have shown in other metabolic pathways—that neutrophils have remarkable metabolic flexibility, which is essential to support their functions as the first responders in innate immunity. Furthermore, the metabolic remodeling in TCA cycle is stimulation-specific. For instance, the strong activation of mitochondrial glucose oxidation occurs with ionomycin but not PMA stimulation. Correlating with this, the impact of mitochondrial perturbation on neutrophil functions is also stimulation-specific. Ionomycin- and MSU crystal-induced NET release is strongly suppressed by MPC inhibitors, but PMA-induced NET release is minimally affected. Such specificity reveals insights on how metabolism is regulated upon different stimulations and how metabolism is mechanistically connected to effector functions. Here, we show calcium influx is necessary for the rapid activation of the TCA cycle in response to certain stimulation (Fig 3B). On the other hand, the differential dependence of NET release on the TCA cycle likely results from the different mechanisms for inducing NET, which pose different demands on redox metabolism [50]. With stimuli that activate NOX-dependent NET release (such as PMA), the activation is primarily enabled by a metabolic switch from glycolysis to a cyclic pentose phosphate pathway to maximize cytosolic NADPH production, which then fuels NOX-dependent ROS production that ultimately induces NET release [17]. With stimulations that do not induce NOX, such as ionomycin, changes in cellular redox are instead driven by an activation of the TCA cycle. ‘Such a context-specific metabolic remodeling and metabolism-function connection can have clinical and therapeutic relevance. TCA cycle remodeling is likely particularly important in contexts that activate calcium signaling, such as MSU stimulation of neutrophils in gouty inflammatory arthritis [51]. Consequentially, perturbation of mitochondrial metabolism would be more relevant for pathologies that involves NOX-independent NET release induced by mitochondrial ROS, such as rheumatoid arthritis, systemic lupus erythematosus, and sepsis [34,52].

The exact mechanism allowing UK-5099 and azemiglitazone to modulate various neutrophil functions requires more investigation. Both drugs can have off-target effects and secondary effects in broader metabolic pathways beyond their shared effect in inhibiting MPC. One notable difference in metabolism is around glycolysis, where azemiglitazone particularly promotes glucose metabolism through glycolysis, while UK-5099 does not (Fig 2D-E). Intriguingly, we also see that while both drugs strongly inhibit NETosis, there are also some different effects. For instance, UK-5099 strongly inhibits neutrophil migration whereas azemiglitazone’s effect on migration is not significant (Fig 5H). Correlating with migration, we also observed different cellular morphology changes. UK-5099 prevented the increase in cell size and eccentricity upon ionomycin stimulation. In contrast, azemiglitazone treatment makes the cells larger and more eccentric (Fig S3). More investigation is needed to understand which specific metabolic perturbations caused by UK-5099 and azemiglitazone are mechanistically linked to which specific functional alterations.

Regardless of the exact mechanism, the fact that these drugs can alter important neutrophil response such as NET release has exciting implications. NETs and MPO release can induce inflammation and tissue damage. Dysregulated NET release is involved in many pathological conditions. The results here point to valuable avenues for manipulating neutrophils in these inflammatory conditions. Azemiglitazone has recently been shown to dampen inflammatory responses of lung macrophages and mitigates hyperinflammation in coronavirus-19 infection [53]. Our study suggests it can act on neutrophils and thus could be useful in neutrophil-driven pathologies as well. The application potential is particularly promising given the safety of azemiglitazone has been evaluated. Azemiglitazone recently passed Phase IIb clinical trials as a treatment for type II diabetes mellitus and metabolic dysfunction associated steatohepatitis [54]. UK-5099 and azemiglitazone are both thiazolidinediones with insulin sensitizing effects [55]. In diabetes, neutrophils are primed for NET release and NETs in diabetic wounds contribute to impaired wound healing [56,57]. From a mechanistic perspective, it is consistent with the idea that altered mitochondrial metabolism, which often occurs in diabetes, can contribute to altered neutrophil response. From the application perspective, the anti-diabetic and anti-NET effects of these inhibitors could provide additional therapeutic benefits to diabetic patients with non-healing wounds. Additional studies to test the anti-NET effects of these inhibitors *in vivo* and their impact on diabetic wounds are merited.

## 4 Figure Legends

**Fig S1.**
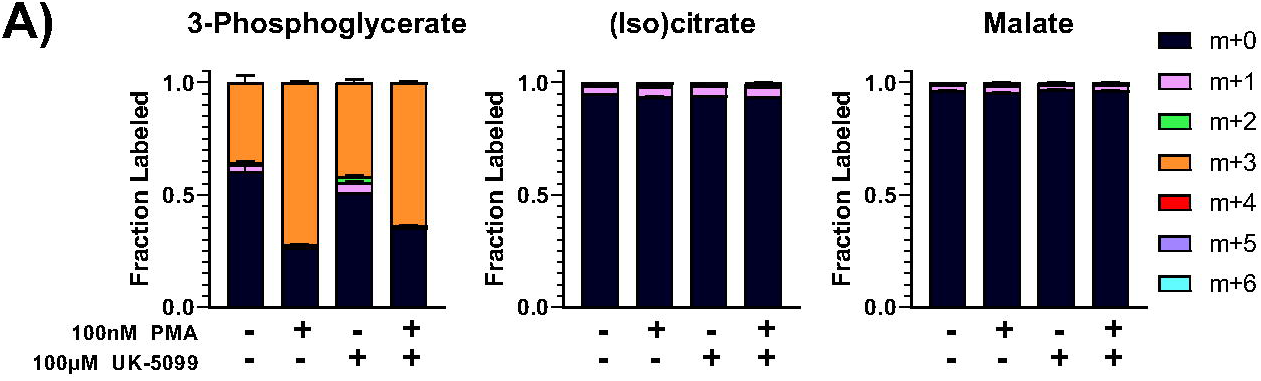
PMA does not increase TCA cycle labeling. U^13^C-glucose labeling into glycolysis and TCA intermediates in primary human neutrophils treated under specified conditions for 1 hour.

**Fig S2.**
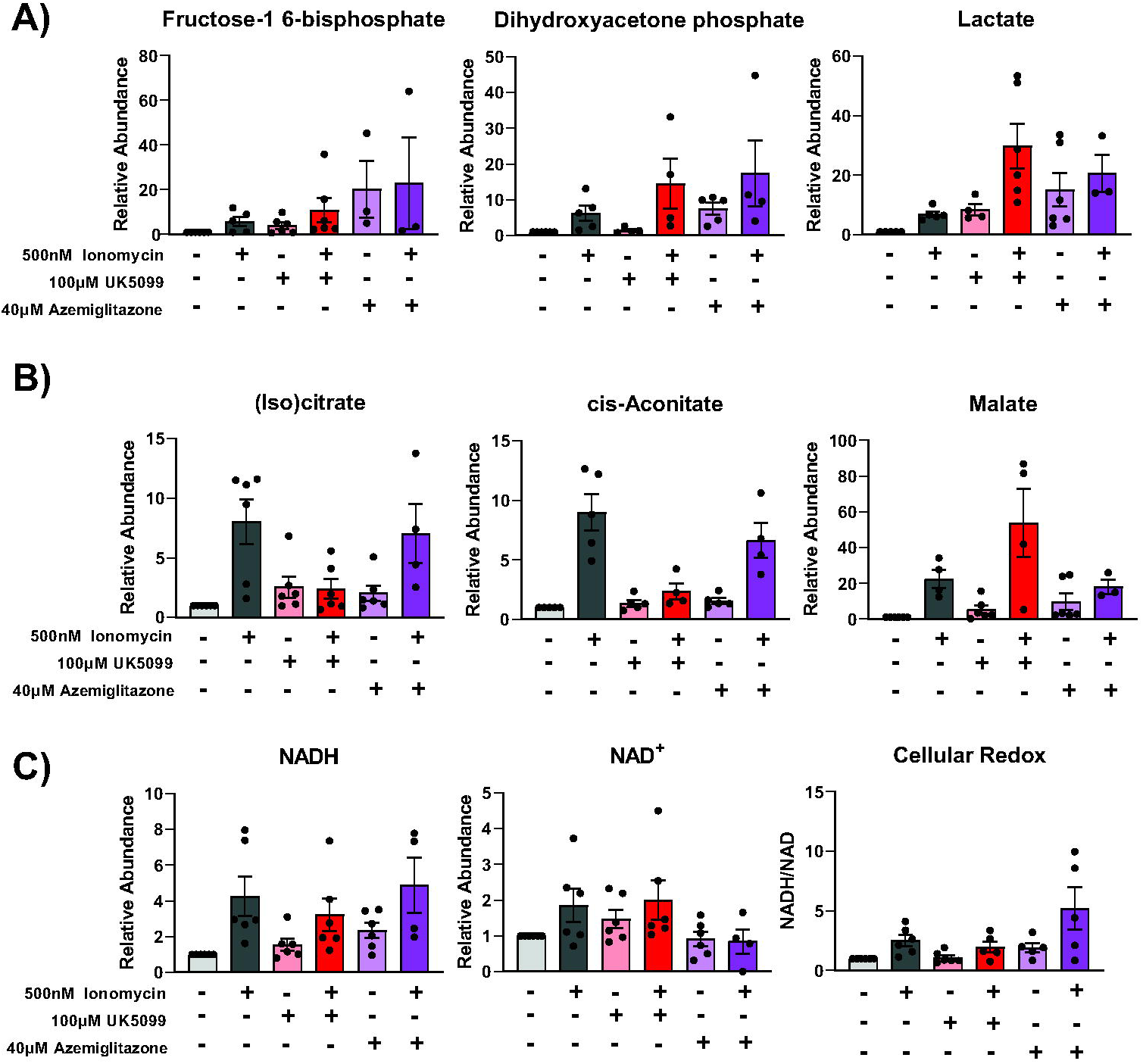
Changes in metabolic abundance in human peripheral blood neutrophils caused by Ionomycin stimulation and MPC inhibition. Relative abundance of (A) glycolysis intermediates, (B) TCA intermediates, and (C) redox metabolites from 6 different donors was compiled and normalized by protein content and made relative to the unstimulated condition from the respective donor. Cells were pretreated with MPC inhibitors for 45 minutes and then stimulated for 1 hour.

**Figure S3.**
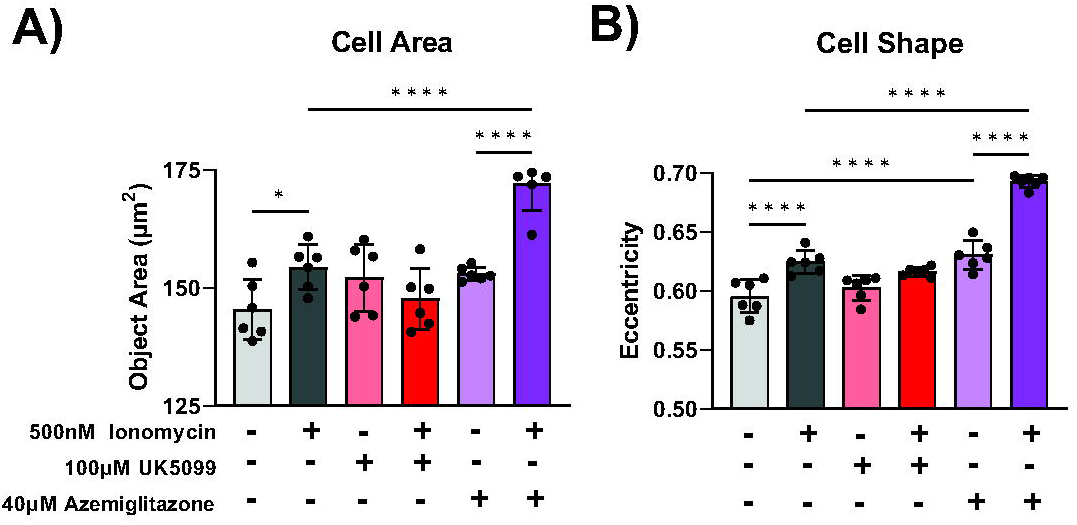
Effects of ionomycin stimulation and MPC inhibition on primary human neutrophil morphology. Cell size (A) and cell eccentricity (B) measurements in neutrophils treated under specified conditions for 4 hours. Significance was determined by ordinary one-way ANOVA and Šídák’s multiple comparisons test.

**Figure S4.**
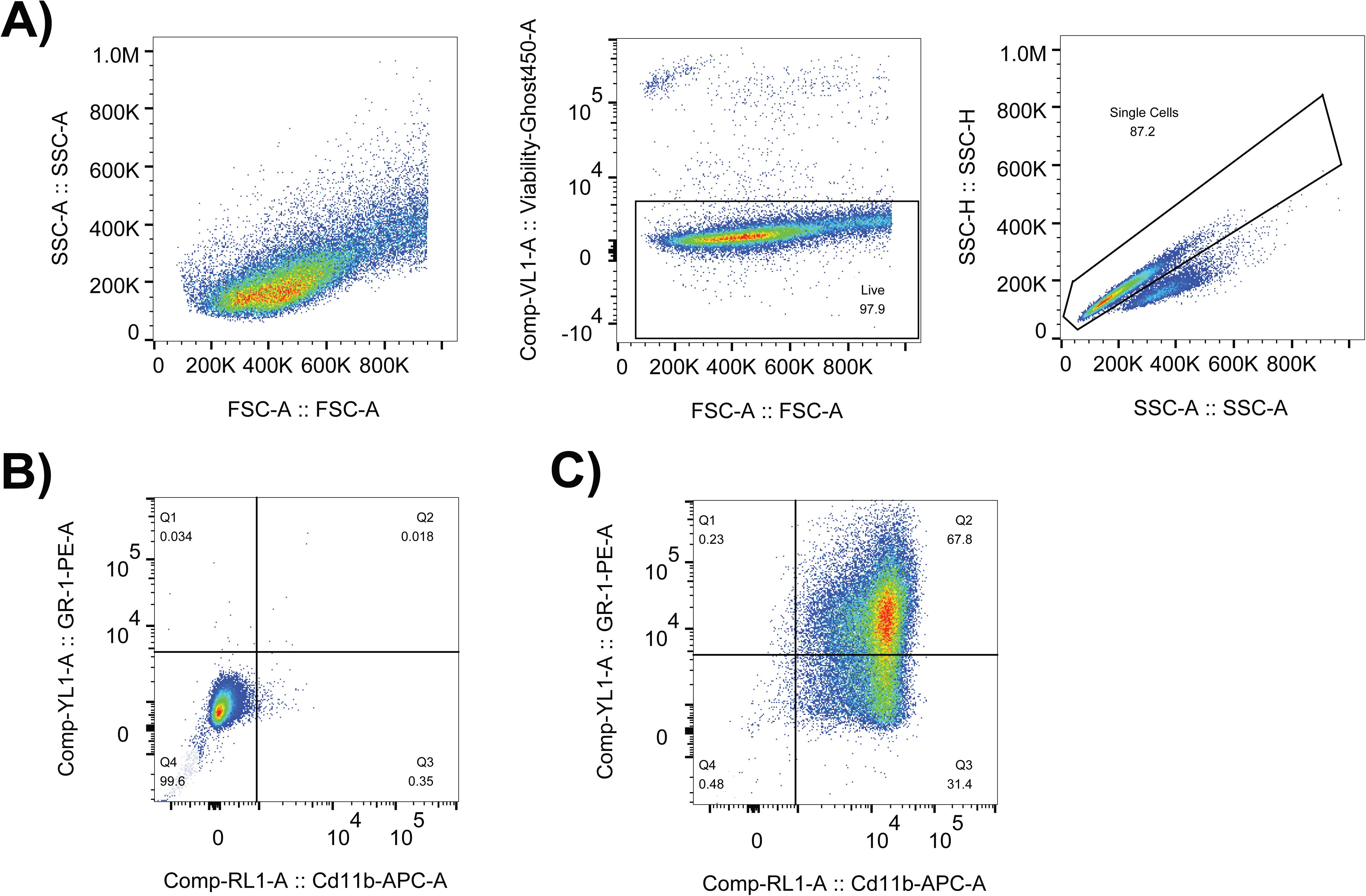
Flow cytometry gating scheme of ER-Hoxb8 neutrophil differentiation. (A) Gating strategy to get live single cells. Gating strategy for CD11b and Gr-1 in (B) undifferentiated and (C) D4 differentiated ER-Hoxb8 neutrophils.

## 5. Materials and Methods

### 5.1 Isolation, culture and stimulation of human peripheral blood neutrophils

Human neutrophils were isolated from peripheral blood freshly collected from healthy donors, following the protocol approved by the University of Wisconsin Institutional Review Board (protocol no. 2019-1031-CP001). Informed consent was obtained from all participants. Neutrophils were purified using the MACSxpress Whole Blood Neutrophil Isolation Kit (Miltenyi Biotec, 130-104-434) followed by erythrocyte depletion (Miltenyi Biotec, 130-098-196) according to the manufacturer’s instructions. Neutrophils were spun down at 300*g* for 5 min, resuspended in neutrophil culture media – RPMI 1640 w/o glucose (ThermoFisher, 11879020) supplemented with 5 mM glucose (Fisher Scientific, D16500) – and kept at 37°C in incubators under 5% CO2. Purity of isolated cells was verified by flow cytometry using antibodies against neutrophil surface markers CD11b-PE (Biolegend, 301305) and CD15-AlexaFluor700 (Biolegend, no. 301919), and cell viability was checked using Ghost Dye 450 (Tonbo Biosciences, 13-0863). Typically, isolated cells are >90% CD11b+ CD15+ and >95% viable as previously reported [17].

### 5.2. Culture and differentiation of murine ER-HOXb8 neutrophils

Cells were cultured at 37°C with 5% CO_2_ in a humidified incubator. ER-Hoxb8 conditionally immortalized myeloid progenitor cells (obtained from the Sykes lab) were cultured and differentiated following previously described protocols [6]. Briefly, undifferentiated ER-Hoxb8 cells were cultured and expanded in RPMI 1640 supplemented with 10% FBS, 100 μg/mL penicillin-streptomycin, 0.5µM β-estradiol, and 2% SCF condition media (from SCF producing CHO cell line also obtained from the Sykes lab). To differentiate the cells, cells were spun down, washed in PBS, and placed in the same media without β-estradiol. For metabolomics throughout differentiation, cells were collected on a specified day, counted and cultured in 1 ml RPMI for 2-hours, washed once in PBS, and metabolites were extracted. Cultured non-differentiating cells were processed as controls on each differentiation day. The differentiation of ER-Hoxb8 neutrophils over time was examined by flow cytometry after staining cells for viability and murine neutrophil surface makers; CD11b (BD Biosciences, 553309) and Gr-1 (BD Biosciences, 553125).

### 5.3. Culture and differentiation of human iPSC-derived neutrophils (iNeutrophils)

iNeutrophils were generated from induced pluripotent stem cells as previously described [22,23]. Briefly, bone marrow-derived human IISH2i-BM9 iPSCs were obtained from WiCell. iPSCs were cultured on Cultrex-coated (WiCell) tissue culture plates in mTeSR-Plus medium (STEMCELL Technologies). To begin the differentiation process, iPSC cells were passaged onto collagen-coated plates (2.4μg/mL) containing TeSR-E8 media with 10μM ROCK inhibitor Y-27632 (Tocris). Cells were then left to adhere to the plate for two hours while incubating at 37°C and 5% CO_2_. Differentiation to hemogenic endothelium (starting at “Day 0”) was initiated via *ETV2* mRNA (TriLink Biotechnologies) transfection in TeSR-E8 media (STEMCELL Technologies) with the use of TransIT reagent and mRNA boost (Mirus Bio). Cells were then left to incubate for one day at 37°C and 5% CO_2_. On day 1, the media was replaced with StemLineII media containing VEGF-165 (20ng/mL; PeproTech) and FGF2 (10ng/mL; PeproTech) to further induce differentiation into hemogenic endothelial cells. This media was then replaced on day 2 with fresh StemLineII media with VEGF-165 and FGF2. At day 3, differentiation from hemogenic endothelia to granulocyte-monocyte progenitors was initiated by changing the media to StemLineII media supplemented with FGF2 (20ng/mL), granulocyte-macrophage colony-stimulating factor (25ng/mL; PeproTech), and UM171 (50nM; Xcess Biosciences). Cells were then left to incubate at 37°C and 5% CO_2_ before topping off the media at day 7 with an equal volume of fresh StemLineII media supplemented FGF2, granulocyte-macrophage colony-stimulating factor and UM171. At day 11, non-adherent progenitor cells were collected and used for iNeutrophil differentiation (these are considered as Day 0 of neutrophil differentiation). Following collection, progenitor cells were cultured in StemSpan SFEM II medium (STEMCELL Technologies), supplemented with GlutaMAX 100× (1×; Thermo Fisher Scientific), ExCyte 0.2% (Merck Millipore), human granulocyte colony-stimulating factor (150ng/mL; PeproTech), and Am580 retinoic acid agonist (2.5μM; STEMCELL Technologies) at 0.8–1 × 10^6^ cells/mL density. After 4 days, suspensions were topped off with equal volumes of StemSpan SFEM II medium supplemented with GlutaMAX, ExCyte, human granulocyte colony-stimulating factor and Am580. iNeutrophils were then collected for metabolomics on days 0, 3 and 7 following progenitor harvesting. Cells were spun down, washed, and metabolites were extracted as described below. By day 7, there is increased expression of neutrophil markers including CD11b, CD15, CD16, CD66b, MPO, and lactoferrin and a gain of effector functions (including chemotaxis, phagocytosis, NETs, and pathogen killing) as previously described [22–24].

### 5.4. Metabolomics and isotopic tracing

Isolated human peripheral blood neutrophils or ER-Hoxb8 neutrophils (2 million/condition) were resuspended into neutrophil culture media and rested for 45 min. In experiments involving inhibitors, cells were pretreated with inhibitors or vehicle control during the 45min period. The cells were then spun down at 500xg for 2 minutes. For metabolomics, cells were resuspended in neutrophil culture media in the presence of inhibitors and/or stimuli as indicated. For isotope tracing, cells were resuspended in RPMI with 5mM U^13^C-glucose (Cambridge Isotope Laboratories, CLM-1396-5) in the place of unlabeled glucose.

To extract intracellular metabolites, culture medium was removed, and neutrophils were immediately washed with PBS. From 2 million pelleted neutrophils, 150 µl of cold liquid chromatography–mass spectrometry (LC–MS)-grade acetonitrile/methanol/water (40:40:20 v:v:v) was added, samples were vortexed and then spun at 20,627*g* for 5 min at 4 C to remove any insoluble debris. Supernatant was then used for LC-MS analysis. Metabolite abundances were normalized by protein content from the same sample. Precipitated protein from metabolite extraction was harvested with RIPA lysis buffer (Thermo Scientific, 89900) with phosphatase inhibitor (Thermo Scientific, A32957) and HALT protease inhibitor (Thermo Scientific, 78425) from the pellet post-metabolite extraction. Total protein concentration was determined by BCA assay kit (ThermoFisher, J63283-QA).

Soluble metabolite samples were analyzed with a Thermo Q-Exactive mass spectrometer coupled to a Vanquish Horizon Ultra-High Performance Liquid Chromatograph, using the following two analytical methods (1). Samples in extraction solvent were directly loaded on to LC–MS, then separated on a 2.1 × 150mm Xbridge BEH Amide (2.5 μm) Column (Waters) using a gradient of solvent A (95% H_2_O, 5% ACN, 20 mM NH_4_AC, 20 mM NH_4_OH) and solvent B (20% H_2_O, 80% ACN, 20 mM NH_4_AC, 20 mM NH_4_OH). The gradient used was 0 min, 100% B; 3 min, 100% B; 3.2 min, 90% B; 6.2 min, 90% B; 6.5 min, 80% B; 10.5 min, 80% B; 10.7 min, 70% B; 13.5 min, 70% B; 13.7 min, 45% B; 16 min, 45% B; 16.5 min, 100% B; 22 min, 100% B. The flow rate was 0.3 ml min^−1^ and column temperature 30 °C. Analytes were measured by MS using full scan (2). Samples were dried under N_2_ flow and resuspended in LC–MS-grade water as loading solvent. Metabolites were separated on a 2.1 × 100mm, 1.7 µM Acquity UPLC BEH C18 Column (Waters) with a gradient of solvent A (97:3 H_2_O/methanol, 10 mM TBA, 9 mM acetate, pH 8.2) and solvent B (100% methanol). The gradient was: 0 min, 5% B; 2.5 min, 5% B; 17 min, 95% B; 21 min, 95% B; 21.5 min, 5% B. Flow rate was 0.2 ml min^−1^. Data were collected with full scan. Identification of metabolites reported here was based on exact *m/z* and retention time that were determined with chemical standards. Data were collected with Xcalibur 4.0 software and analyzed with Maven. Quantitative Enrichment Analysis was performed using MetaboAnalyst 6.0.

To determine energy charge, relative metabolite abundance of AMP, ADP, and ATP was quantified from ion count in LC-MS analysis based on external calibration curves obtained by measuring a series of AMP, ADP, and ATP standards at various concentration using the same LC-MS method. Energy charge is calculated based on Equation 1.

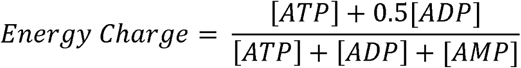

### 5.5. Quantification of oxygen consumption rate (OCR)

OCR was measured using XF-96e extracellular flux analyzer (Agilent). To attach neutrophils at the bottom of assay plates (Seahorse Bioscience), neutrophils were plated in culture wells precoated with Cell-Tak (Corning, no. 354240) at 4.5 × 10^5^ cells per well, spun at 100g for 1 min with minimal acceleration/deceleration then incubated for 1 h at 37 °C. Assays were performed in neutrophil culture media.

### 5.6. Sample preparation for multiphoton imaging

Neutrophils were plated at 5 × 10^4^ cells per well in a 96-well glass bottom plate (Cellvis, Catalog no. P96-1.5H-N) precoated with Cell-Tak (Corning, no. 354240) and spun at 100*g* for 1 min with minimal acceleration/deceleration. Cells were then pretreated in inhibitor or vehicle control. Then, neutrophils were stimulated 90 minutes before imaging. During imaging, the samples were housed in a stage top incubator (Tokai Hit) to keep them at 37°C and under 5% CO_2_.

### 5.7. Optical metabolic imaging

Two-photon fluorescence-lifetime imaging microscopy (FLIM) was performed on a custom-made Ultima Multiphoton Imaging System (Bruker) that consists of an inverted microscope (TI-E, Nikon). The system is coupled to an ultrafast tunable laser source (Insight DS+, Spectra Physics Inc). The fluorescence lifetime images were acquired using time-correlated single-photon counting (TSCPC) electronics (SPC -150, Becker & Hickl GmbH) and imaging was performed using Prairie View Software (Bruker). NAD(P)H and FAD were sequentially excited using excitation wavelengths of 750 nm and 890 nm respectively, and the laser power at the sample was <10 mW. The samples were illuminated using a 40x objective lens (W.I./1.15 NA /Nikon PlanApo) with a pixel dwell time of 4.8 µs and frame integration of 60 s at 256×256 pixels and 2x zoom of maximum field of view (∼0.9 mm^2^). The photon count rates were maintained at 1–5 × 10^5^ photons/second to ensure adequate photon detection for accurate lifetime decay fits. A dichroic mirror (720 nm) and bandpass filters separated fluorescence signals from the excitation laser. Emission filters used were bandpass 460/80 nm for NAD(P)H and 500/100 nm for FAD. Fluorescence signals were collected on GaAsP photomultiplier tubes (H7422P-40, Hamamatsu, Japan). The instrument response function (IRF) was collected each day by recording the second harmonic generation signal of urea crystals (Sigma-Aldrich) excited at 890nm.

### 5.8. Image analysis and single-cell quantification

Fluorescence lifetime images were analyzed on SPCImage software (Becker & Hickl). Fluorescence counts at each pixel were enhanced by a 3x3 binning comprising of 9 surrounding pixels. To eliminate pixels with a low fluorescence signal, an intensity threshold was used. NAD(P)H has a closed conformation in its bound state while it adopts an open conformation in free state and hence has a comparatively shorter fluorescence lifetime. Conversely, FAD has a long lifetime in the free state and a shorter lifetime in bound state. Thus, using an iterative parameter optimization to obtain the lowest sum of the squared differences between model and data (Weighted Least Squares algorithm), the pixel-wise NAD(P)H and FAD decay curves were fit to a biexponential model [I(t) = α_1_×exp (−t/t_1_) + α_2_×exp(−t/t_2_) + C)] convolved with the system IRF. Here, I(t) represent the fluorescence intensity measured at time t, α_1_, α_2_ are the fractional contributions and t_1_, t_2_ denote the short and long lifetime components, respectively. C accounts for background light. The goodness of the fit was checked using a reduced chi-squared value<1.0. To generate the intensity of NAD(P)H (I_NAD(P)H_) and FAD (I_FAD_), the area under the fluorescence decay curve at each pixel was integrated. The optical redox ratio at each pixel is given by [I_NAD(P)H_ / (_INAD(P)H_ + I_FAD_)].

Automated whole cell masks were created from NAD(P)H intensity images using Cellpose 2.0 (model = cyto 2, radius = 30) [25]. Following this, the generated masks were manually edited on napari [26]. All the processing and analysis of single-cell data was done using a custom python library Cell Analysis Tools that extracted OMI and morphological features for each cell mask [27].

### 5.9. NET release assay

NET release was quantified by the increase in extracellular DNA over time following a previously developed protocol (PMID 29196457). Briefly, neutrophils were plated at 5 × 10^4^ cells per well in a 96-well tissue culture plate precoated with Cell-Tak (Corning, no. 354240) and spun at 100*g* for 1 min with minimal acceleration/deceleration. Cytotox Green Reagent (Sartorius, 4633) was added to culture media at a 1:4,000 dilution to stain extracellular DNA, and images were captured every 20–60 min after stimulation using an IncuCyte live cell imager (maker) under standard culture conditions (37 °C, 5% CO_2_). The fluorescent area outside of cells, which indicates NETs, was quantified by image analysis using IncuCyte S3 Basic Analysis software.

### 5.10. MPO release assay

Myeloperoxidase release was assayed using a Neutrophil Myeloperoxidase Activity Assay Kit (Cayman Chemical, catalog no. 600620) following manufacturer instructions. Briefly, neutrophils were plated at 2.5 × 10^5^ cells per well in a 96-well tissue culture plate, pretreated for 45 minutes with inhibitor or vehicle control, and stimulated for 4 hours. Plate was then spun down at 1000*g* for 8min and supernatant was collected and analyzed for MPO release.

### 5.11. Immunoblotting

1 million cells were lysed with Laemmli lysis buffer with phosphatase inhibitor (Thermo Scientific, no. A32957) and HALT protease inhibitor (Thermo Scientific, 78425). Proteins of interest were probed with the following antibodies in TBS-T buffer with 5% BSA. Primary antibodies: Citrullinated Histone H3 (Cell Signaling, E4O3F), Histone H3 (Cell Signaling, #9715), phospho-PDH [ser293] (Novus Biologicals, NB110-93479), phospho-PDH [ser300] (Millipore Sigma, ABS194), PDH (Cell Signaling, #2784), β-actin (Cell Signaling, 4967); and secondary antibodies: Goat-anti-rabbit 800 (LI-COR, 925-32211), Goat-antimouse 680 (LI-COR, 925-68070). Membranes were imaged with a Li-Cor Odyssey ClX and quantified using ImageStudio Lite software.

### 5.12. Neutrophil migration assay

Migration assays were performed across a C5a gradient through transwell inserts with 3 µm pores (Corning 3415). Isolated primary neutrophils were rested for 30 min in RPMI 1640 medium containing 11mM glucose and 0.1% HSA, then pre-treated for 30 min with experimental condition. Then 2×10^5^ cells were placed into each loading control wells or transwell and migrated towards 2 nM C5a (biotechne, 2037-C5) added to the other side of transwell. After 1.5 hours, transwells were removed and images were taken of the bottom well containing the migrated cells. Cells were quantified using the IncuCyte S3 Basic Analysis software. Cell loading was checked for every condition by quantifying the cells in loading control wells under the same conditions.

### 5.13. Apoptosis assay

Apoptosis was quantified by the fraction of cells stained positive for AnnexinV red (Sartorius, 4641) and negative for Cytotox Green over time. Cell culture and imaging plate was set up in a similar procedure described above for NET release assay. In addition to Cytotox green dye, AnnexinV red dye was added at a 1:200 dilution (per manufacturer protocol). Total cells and cells staining positive for AnnexinV, Cytotox, or both were quantified and analyzed using IncuCyte S3 adherent cell-by-cell analysis software. For cell morphology measurements, images were taken using the 20x objective and IncuCyte S3 non-adherent cell-by-cell analysis was used for cell masking.

### 5.14. Quantification and statistical analysis

Data are presented as mean ± SD for technical replicates or mean ± SEM for biological replicates. Outliers were identified by ROUT (Q = 1%) and excluded from analysis. Statistical significance is indicated in all figures by the following annotations: *p < 0.05, **p < 0.01, ***p < 0.001, ****p < 0.0001.

## 6 Acknowledgements

The authors thank the University of Wisconsin Carbone Cancer Center Flow Cytometry Laboratory, supported by P30 CA014520, for use of its facilities and services. Training grant T32GM140935 from the National Institute of General Medical Sciences (https://www.nigms.nih.gov/) supports JL. Training grants TL1TR002375 and UL1TR002373 from the National Center for Advancing Translational Sciences (https://ncats.nih.gov/) support JL. Grant R21AI5229 was received from the National Institute of Allergy and Infectious Diseases (https://www.niaid.nih.gov/) to fund AH, Grant U24AI152177 was received from the National Institute of Allergy and Infectious Diseases (https://www.niaid.nih.gov/) to fund MS. Grant R35GM147014 was received from the National Institute of General Medical Sciences (https://www.nigms.nih.gov/) to fund JF.

## 7. Declaration of Competing Interest

The authors declare that they have no known competing financial interests or personal relationships that could have appeared to influence the work reported in this paper.

## Notes

### Competing Interest Statement

The authors have declared no competing interest.

## References

[1] Mann, M., Mehta, A., de Boer, C.G., Kowalczyk, M.S., Lee, K., Haldeman, P., et al., 2018. Heterogeneous Responses of Hematopoietic Stem Cells to Inflammatory Stimuli Are Altered with Age. Cell Reports 25(11): 2992–3005.e5, Doi: 10.1016/j.celrep.2018.11.056.

[2] Kwok, I., Becht, E., Xia, Y., Ng, M., Teh, Y.C., Tan, L., et al., 2020. Combinatorial Single-Cell Analyses of Granulocyte-Monocyte Progenitor Heterogeneity Reveals an Early Uni-potent Neutrophil Progenitor. Immunity 53(2): 303–318.e5, Doi: 10.1016/j.immuni.2020.06.005.

[3] Thind, M.K., Uhlig, H.H., Glogauer, M., Palaniyar, N., Bourdon, C., Gwela, A., et al., 2023. A metabolic perspective of the neutrophil life cycle: new avenues in immunometabolism. Frontiers in Immunology 14: 1334205, Doi: 10.3389/fimmu.2023.1334205.

[4] Koenderman, L., Tesselaar, K., Vrisekoop, N., 2022. Human neutrophil kinetics: a call to revisit old evidence. Trends in Immunology 43(11): 868–76, Doi: 10.1016/j.it.2022.09.008.

[5] Hong, C.-W., 2017. Current Understanding in Neutrophil Differentiation and Heterogeneity. Immune Network 17(5): 298–306, Doi: 10.4110/in.2017.17.5.298.

[6] Wang, G.G., Calvo, K.R., Pasillas, M.P., Sykes, D.B., Häcker, H., Kamps, M.P., 2006. Quantitative production of macrophages or neutrophils ex vivo using conditional Hoxb8. Nature Methods 3(4): 287–93, Doi: 10.1038/nmeth865.

[7] Jafarzadeh, A., Motaghi, M., Patra, S.K., Jafarzadeh, Z., Nemati, M., Saha, B., 2024. Neutrophil generation from hematopoietic progenitor cells and induced pluripotent stem cells (iPSCs): potential applications. Cytotherapy 26(8): 797–805, Doi: 10.1016/j.jcyt.2024.03.483.

[8] Petri, B., Sanz, M.-J., 2018. Neutrophil chemotaxis. Cell and Tissue Research 371(3): 425– 36, Doi: 10.1007/s00441-017-2776-8.

[9] Manfredi, A.A., Ramirez, G.A., Rovere-Querini, P., Maugeri, N., 2018. The Neutrophil’s Choice: Phagocytose vs Make Neutrophil Extracellular Traps. Frontiers in Immunology 9: 288, Doi: 10.3389/fimmu.2018.00288.

[10] Mutua, V., Gershwin, L.J., 2021. A Review of Neutrophil Extracellular Traps (NETs) in Disease: Potential Anti-NETs Therapeutics. Clinical Reviews in Allergy & Immunology 61(2): 194–211, Doi: 10.1007/s12016-020-08804-7.

[11] Mayadas, T.N., Cullere, X., Lowell, C.A., 2014. The Multifaceted Functions of Neutrophils. Annual Review of Pathology: Mechanisms of Disease 9(1): 181–218, Doi: 10.1146/annurev-pathol-020712-164023.

[12] Lika, J., Fan, J., 2024. Carbohydrate metabolism in supporting and regulating neutrophil effector functions. Current Opinion in Immunology 91: 102497, Doi: 10.1016/j.coi.2024.102497.

[13] Arnold, P.K., Finley, L.W.S., 2023. Regulation and function of the mammalian tricarboxylic acid cycle. Journal of Biological Chemistry 299(2): 102838, Doi: 10.1016/j.jbc.2022.102838.

[14] Wang, M., Zhu, B., Zhang, C., Li, C., Zhang, R., Rathmell, J., et al., 2024. Glutamine Metabolism Is Required for Alveolar Macrophage Proliferation. Journal of Respiratory Biology and Translational Medicine 1(1): 10004–10004, Doi: 10.35534/jrbtm.2024.10004.

[15] Krysa, S.J., Allen, L.-A.H., 2022. Metabolic Reprogramming Mediates Delayed Apoptosis of Human Neutrophils Infected With Francisella tularensis. Frontiers in Immunology 13: 836754, Doi: 10.3389/fimmu.2022.836754.

[16] Cluntun, A.A., Badolia, R., Lettlova, S., Parnell, K.M., Shankar, T.S., Diakos, N.A., et al., 2021. The pyruvate-lactate axis modulates cardiac hypertrophy and heart failure. Cell Metabolism 33(3): 629–648.e10, Doi: 10.1016/j.cmet.2020.12.003.

[17] Britt, E.C., Lika, J., Giese, M.A., Schoen, T.J., Seim, G.L., Huang, Z., et al., 2022. Switching to the cyclic pentose phosphate pathway powers the oxidative burst in activated neutrophils. Nature Metabolism 4(3): 389–403, Doi: 10.1038/s42255-022-00550-8.

[18] Sadiku, P., Willson, J.A., Ryan, E.M., Sammut, D., Coelho, P., Watts, E.R., et al., 2021. Neutrophils Fuel Effective Immune Responses through Gluconeogenesis and Glycogenesis. Cell Metabolism 33(2): 411–423.e4, Doi: 10.1016/j.cmet.2020.11.016.

[19] Kumar, S., Dikshit, M., 2019. Metabolic Insight of Neutrophils in Health and Disease. Frontiers in Immunology 10: 2099, Doi: 10.3389/fimmu.2019.02099.

[20] Forrest, O.A., Ingersoll, S.A., Preininger, M.K., Laval, J., Limoli, D.H., Brown, M.R., et al., 2018. Frontline Science: Pathological conditioning of human neutrophils recruited to the airway milieu in cystic fibrosis. Journal of Leukocyte Biology 104(4): 665–75, Doi: 10.1002/JLB.5HI1117-454RR.

[21] Cao, Z., Zhao, M., Sun, H., Hu, L., Chen, Y., Fan, Z., 2022. Roles of mitochondria in neutrophils. Frontiers in Immunology 13: 934444, Doi: 10.3389/fimmu.2022.934444.

[22] Giese, M.A., Bennin, D.A., Schoen, T.J., Peterson, A.N., Schrope, J.H., Brand, J., et al., 2024. PTP1B phosphatase dampens iPSC-derived neutrophil motility and antimicrobial function. Journal of Leukocyte Biology 116(1): 118–31, Doi: 10.1093/jleuko/qiae039.

[23] Majumder, A., Suknuntha, K., Bennin, D., Klemm, L., Brok-Volchanskaya, V.S., Huttenlocher, A., et al., 2020. Generation of Human Neutrophils from Induced Pluripotent Stem Cells in Chemically Defined Conditions Using ETV2 Modified mRNA. STAR Protocols 1(2): 100075, Doi: 10.1016/j.xpro.2020.100075.

[24] Wagner, A.S., Smith, F.M., Bennin, D.A., Votava, J.A., Datta, R., Giese, M.A., et al., 2024. GATA1-deficient human pluripotent stem cells generate neutrophils with improved antifungal immunity that is mediated by the integrin CD18. bioRxiv: The Preprint Server for Biology: 2024.10.11.617742, Doi: 10.1101/2024.10.11.617742.

[25] Pachitariu, M., Stringer, C., 2022. Cellpose 2.0: how to train your own model. Nature Methods 19(12): 1634–41, Doi: 10.1038/s41592-022-01663-4.

[26] Sofroniew, N., Lambert, T., Evans, K., Nunez-Iglesias, J., Bokota, G., Winston, P., et al., 2022. napari: a multi-dimensional image viewer for Python, Doi: 10.5281/ZENODO.7276432.

[27] Contreras Guzman, E., Rehani, P., Skala, M.C., 2023. Cell analysis tools: an open-source library for single-cell analysis of multi-dimensional microscopy images. In: Tarnok, A., Houston, J.P., Su, X., editors. Imaging, Manipulation, and Analysis of Biomolecules, Cells, and Tissues XXI, San Francisco, United States: SPIE p. 31.

[28] Liang, R., Arif, T., Kalmykova, S., Kasianov, A., Lin, M., Menon, V., et al., 2020. Restraining Lysosomal Activity Preserves Hematopoietic Stem Cell Quiescence and Potency. Cell Stem Cell 26(3): 359–376.e7, Doi: 10.1016/j.stem.2020.01.013.

[29] Wang, L., Ai, Z., Khoyratty, T., Zec, K., Eames, H.L., van Grinsven, E., et al., 2020. ROS-producing immature neutrophils in giant cell arteritis are linked to vascular pathologies. JCI Insight 5(20): e139163, Doi: 10.1172/jci.insight.139163.

[30] Britt, E.C., Qing, X., Votava, J.A., Lika, J., Wagner, A.S., Shen, S., et al., 2024. Activation induces shift in nutrient utilization that differentially impacts cell functions in human neutrophils. Proceedings of the National Academy of Sciences 121(39): e2321212121, Doi: 10.1073/pnas.2321212121.

[31] Benova, A., Ferencakova, M., Bardova, K., Funda, J., Prochazka, J., Spoutil, F., et al., 2022. Novel thiazolidinedione analog reduces a negative impact on bone and mesenchymal stem cell properties in obese mice compared to classical thiazolidinediones. Molecular Metabolism 65: 101598, Doi: 10.1016/j.molmet.2022.101598.

[32] Yeaman, S.J., Hutcheson, E.T., Roche, T.E., Pettit, F.H., Brown, J.R., Reed, L.J., et al., 1978. Sites of phosphorylation on pyruvate dehydrogenase from bovine kidney and heart. Biochemistry 17(12): 2364–70, Doi: 10.1021/bi00605a017.

[33] Denton, R.M., 2009. Regulation of mitochondrial dehydrogenases by calcium ions. Biochimica Et Biophysica Acta 1787(11): 1309–16, Doi: 10.1016/j.bbabio.2009.01.005.

[34] Douda, D.N., Khan, M.A., Grasemann, H., Palaniyar, N., 2015. SK3 channel and mitochondrial ROS mediate NADPH oxidase-independent NETosis induced by calcium influx. Proceedings of the National Academy of Sciences 112(9): 2817–22, Doi: 10.1073/pnas.1414055112.

[35] Naccache, P.H., Grimard, M., Roberge, C.J., Gilbert, C., Poubelle, P.E., Lussier, A., et al., 1991. Crystal induced neutrophil activation. I. Initiation and modulation of calcium mobilization and superoxide production by microcrystals. Arthritis & Rheumatism 34(3): 333–42, Doi: 10.1002/art.1780340311.

[36] Gupta, A.K., Giaglis, S., Hasler, P., Hahn, S., 2014. Efficient Neutrophil Extracellular Trap Induction Requires Mobilization of Both Intracellular and Extracellular Calcium Pools and Is Modulated by Cyclosporine A. PLoS ONE 9(5): e97088, Doi: 10.1371/journal.pone.0097088.

[37] Datta, R., Miskolci, V., Gallego-López, G.M., Britt, E., Gillette, A., Kralovec, A., et al., 2024. Single cell autofluorescence imaging reveals immediate metabolic shifts of neutrophils with activation across biological systems. bioRxiv: The Preprint Server for Biology: 2024.07.26.605362, Doi: 10.1101/2024.07.26.605362.

[38] Parker, H., Dragunow, M., Hampton, M.B., Kettle, A.J., Winterbourn, C.C., 2012. Requirements for NADPH oxidase and myeloperoxidase in neutrophil extracellular trap formation differ depending on the stimulus. Journal of Leukocyte Biology 92(4): 841–9, Doi: 10.1189/jlb.1211601.

[39] Beerman, R.W., Matty, M.A., Au, G.G., Looger, L.L., Choudhury, K.R., Keller, P.J., et al., 2015. Direct In Vivo Manipulation and Imaging of Calcium Transients in Neutrophils Identify a Critical Role for Leading-Edge Calcium Flux. Cell Reports 13(10): 2107–17, Doi: 10.1016/j.celrep.2015.11.010.

[40] Bao, Y., Ledderose, C., Seier, T., Graf, A.F., Brix, B., Chong, E., et al., 2014. Mitochondria regulate neutrophil activation by generating ATP for autocrine purinergic signaling. The Journal of Biological Chemistry 289(39): 26794–803, Doi: 10.1074/jbc.M114.572495.

[41] Hoogendijk, A.J., Pourfarzad, F., Aarts, C.E.M., Tool, A.T.J., Hiemstra, I.H., Grassi, L., et al., 2019. Dynamic Transcriptome-Proteome Correlation Networks Reveal Human Myeloid Differentiation and Neutrophil-Specific Programming. Cell Reports 29(8): 2505–2519.e4, Doi: 10.1016/j.celrep.2019.10.082.

[42] Maianski, N.A., Geissler, J., Srinivasula, S.M., Alnemri, E.S., Roos, D., Kuijpers, T.W., 2004. Functional characterization of mitochondria in neutrophils: a role restricted to apoptosis. Cell Death & Differentiation 11(2): 143–53, Doi: 10.1038/sj.cdd.4401320.

[43] Reiss, M., Roos, D., 1978. Differences in Oxygen Metabolism of Phagocytosing Monocytes and Neutrophils. Journal of Clinical Investigation 61(2): 480–8, Doi: 10.1172/JCI108959.

[44] Liu, S., Wu, W., Du, Y., Yin, H., Chen, Q., Yu, W., et al., 2023. The evolution and heterogeneity of neutrophils in cancers: origins, subsets, functions, orchestrations and clinical applications. Molecular Cancer 22(1): 148, Doi: 10.1186/s12943-023-01843-6.

[45] Bissenova, S., Ellis, D., Mathieu, C., Gysemans, C., 2022. Neutrophils in autoimmunity: when the hero becomes the villain. Clinical and Experimental Immunology 210(2): 128–40, Doi: 10.1093/cei/uxac093.

[46] Rice, C.M., Davies, L.C., Subleski, J.J., Maio, N., Gonzalez-Cotto, M., Andrews, C., et al., 2018. Tumour-elicited neutrophils engage mitochondrial metabolism to circumvent nutrient limitations and maintain immune suppression. Nature Communications 9(1): 5099, Doi: 10.1038/s41467-018-07505-2.

[47] Britt, E.C., John, S.V., Locasale, J.W., Fan, J., 2020. Metabolic regulation of epigenetic remodeling in immune cells. Current Opinion in Biotechnology 63: 111–7, Doi: 10.1016/j.copbio.2019.12.008.

[48] Martínez-Reyes, I., Chandel, N.S., 2020. Mitochondrial TCA cycle metabolites control physiology and disease. Nature Communications 11(1): 102, Doi: 10.1038/s41467-019-13668-3.

[49] Williams, N.C., O’Neill, L.A.J., 2018. A Role for the Krebs Cycle Intermediate Citrate in Metabolic Reprogramming in Innate Immunity and Inflammation. Frontiers in Immunology 9: 141, Doi: 10.3389/fimmu.2018.00141.

[50] Kenny, E.F., Herzig, A., Krüger, R., Muth, A., Mondal, S., Thompson, P.R., et al., 2017. Diverse stimuli engage different neutrophil extracellular trap pathways. eLife 6: e24437, Doi: 10.7554/eLife.24437.

[51] Cao, X., Li, Y., Luo, Y., Chu, T., Yang, H., Wen, J., et al., 2023. Transient receptor potential melastatin 2 regulates neutrophil extracellular traps formation and delays resolution of neutrophil-driven sterile inflammation. Journal of Inflammation 20(1): 7, Doi: 10.1186/s12950-023-00334-1.

[52] Pieterse, E., Rother, N., Yanginlar, C., Gerretsen, J., Boeltz, S., Munoz, L.E., et al., 2018. Cleaved N-terminal histone tails distinguish between NADPH oxidase (NOX)-dependent and NOX-independent pathways of neutrophil extracellular trap formation. Annals of the Rheumatic Diseases 77(12): 1790–8, Doi: 10.1136/annrheumdis-2018-213223.

[53] Zhu, B., Wei, X., Narasimhan, H., Qian, W., Zhang, R., Cheon, I.S., et al., 2023. Inhibition of the mitochondrial pyruvate carrier simultaneously mitigates hyperinflammation and hyperglycemia in COVID-19. Science Immunology 8(82): eadf0348, Doi: 10.1126/sciimmunol.adf0348.

[54] Harrison, S.A., Alkhouri, N., Davison, B.A., Sanyal, A., Edwards, C., Colca, J.R., et al., 2020. Insulin sensitizer MSDC-0602K in non-alcoholic steatohepatitis: A randomized, double-blind, placebo-controlled phase IIb study. Journal of Hepatology 72(4): 613–26, Doi: 10.1016/j.jhep.2019.10.023.

[55] Divakaruni, A.S., Wiley, S.E., Rogers, G.W., Andreyev, A.Y., Petrosyan, S., Loviscach, M., et al., 2013. Thiazolidinediones are acute, specific inhibitors of the mitochondrial pyruvate carrier. Proceedings of the National Academy of Sciences 110(14): 5422–7, Doi: 10.1073/pnas.1303360110.

[56] Wong, S.L., Demers, M., Martinod, K., Gallant, M., Wang, Y., Goldfine, A.B., et al., 2015. Diabetes primes neutrophils to undergo NETosis, which impairs wound healing. Nature Medicine 21(7): 815–9, Doi: 10.1038/nm.3887.

[57] Shrestha, S., Lee, Y.-B., Lee, H., Choi, Y.-K., Park, B.-Y., Kim, M.-J., et al., 2024. Diabetes Primes Neutrophils for Neutrophil Extracellular Trap Formation through Trained Immunity. Research (Washington, D.C.) 7: 0365, Doi: 10.34133/research.0365.

